# NADK upregulation is an essential metabolic adaptation that enables breast cancer metastatic colonization

**DOI:** 10.1101/2022.03.25.485887

**Authors:** Didem Ilter, Tanya Schild, Nathan P. Ward, Stanislav Drapela, Emma Adhikari, Vivien Low, Eric Le-Lau, Gina M. DeNicola, Melanie R. McReynolds, Ana P. Gomes

## Abstract

Metabolic reprogramming and metabolic plasticity allow cancer cells to fine-tune their metabolism to adapt to the ever-changing environments of the metastatic cascade, for which lipid metabolism and oxidative stress are of particular importance. Continuous production of NADPH, a cornerstone of both lipid and redox homeostasis, is essential for proliferation and cell survival suggesting that cancer cells may require larger pools of NADPH to efficiently metastasize. NADPH is recycled through reduction of NADP+ by several enzymatic systems in cells; however, *de novo* NADP+ is synthesized only through one known enzymatic reaction, catalysed by NAD+ kinase (NADK). Here, we show that NADK is upregulated in metastatic breast cancer cells enabling *de novo* production of NADP(H) and the expansion of the NADP(H) pools thereby increasing the ability of these cells to adapt to the oxidative challenges of the metastatic cascade and efficiently metastasize. Mechanistically, we found that metastatic signals lead to a histone H3.3 variant-mediated epigenetic regulation of the NADK promoter, resulting in increased NADK levels in cells with metastatic ability. Together, our work presents a previously uncharacterized role for NADK as an important contributor to breast cancer progression and suggests that NADK constitutes an important and much needed therapeutic target for metastatic breast cancers.

## Main Text

Breast cancer is the most common malignancy that affects Western women. While the primary tumour can be treated by surgery and adjuvant therapy, metastases are a major cause of mortality (1-3). Thus, the ability to effectively treat solid tumours is largely dependent on the capacity to prevent and/or treat metastatic disease. Metastasis is a complex process in which cancer cells escape the primary tumour, invade the adjacent tissue, enter into and survive in circulation and eventually colonize distant sites (1-3). As such, cancer cells that go through the metastatic cascade need to overcome many obstacles and adapt to many different environments to retain vitality and eventually thrive as metastatic lesions. Metabolic reprogramming and metabolic plasticity provide cancer cells with the flexibility necessary to withstand these challenges and power the development of metastatic disease (4, 5). Lipid metabolism and oxidative stress are of particular importance in the metastatic cascade (4, 6-9). In fact, disruption of either lipid or redox homeostasis has profound effects in the ability of cancer cells to thrive as metastases (10-14).

A cornerstone of both lipid and redox homeostasis is nicotinamide adenine dinucleotide phosphate (NADP+) metabolism. Reduced NADP (NADPH) powers biosynthetic pathways (including lipid synthesis) as well as redox homeostasis (15) and continuous production of NADPH is essential for cell survival and proliferation (16). Thus, we questioned if maintaining NADPH levels is important for breast cancer metastasis. To test this, we expressed triphosphopyridine nucleotide oxidase (TPNOX) in the highly metastatic 4T1 triple negative breast cancer (TNBC) model. TPNOX is an engineered mutant of the naturally occurring *Lactobacillus brevis* NADH oxidase that is strictly specific towards NADPH and whose expression leads to NADPH oxidation into NADP+ thereby decreasing NADPH availability (17). To evaluate the effects of TPNOX expression on the metastatic ability of 4T1 cells, we grew these cells in 3D conditions, which have been shown to be a better surrogate than 2D growth for the metabolic alterations that occur in metastasis (18). TPNOX expression severely blunted the ability of these cells to grow in soft agar (Fig. 1a), demonstrating the importance of NADPH for the 3D growth and metastatic potential of these cells. Based on these results, we hypothesized that breast cancer cells may hijack pathways that reduce NADP+ into NADPH to maintain the NADPH levels necessary for 3D growth and efficient metastasis. To test this we took advantage of clonal subpopulations, which were isolated from a 4T1 mammary tumor and display different metastatic abilities (19), to measure NADP+ and NADPH levels. NADPH levels were significantly higher in the broadly metastatic 4T1 clone, which can form metastases, compared to a locally invasive 4T07 clone, which can invade out of the primary tumor but fails to form metastases (Fig. 1b). Unexpectedly, we observed no change in NADP+ levels (Fig. 1c) putting forward the idea that breast cancer cells with metastatic ability may increase their total NADP(H) pools. Changes in the total NADP(H) pools are regulated by *de novo* NADP(H) synthesis, via phosphorylation of nicotinamide adenine dinucleotide (NAD+) to form NADP+, which is catalyzed by NAD kinases (NADKs) (15) (Fig. 1d). To evaluate if indeed metastatic proficient cells have higher rates of *de novo* NADP(H) synthesis we traced the incorporation of isotopic label from deuterated nicotinamide, [^2^H]NAM, into the NADP(H) pools (20). We observed increased label incorporation into the NADP+ and NADPH pools of the 4T1 clone when compared to the 4T07 clone (Fig. 1e and 1f), demonstrating that *de novo* NADP(H) synthesis is increased in breast cancer cells with the ability to effectively metastasize. Mammalian *de novo* NADP(H) synthesis can be catalyzed by two NADK isoforms, a cytosolic (NADK) and a mitochondrial one (NADK2) (15). Considering that the increase in reactive oxygen species (ROS) in disseminated cancer cells has previously been shown to occur in the cytosol before it propagates to both cytosol and mitochondria in metastatic lesions (10) as well as the fact that fatty acid synthesis occurs in the cytosol, we reasoned that the cytosolic NADK might be responsible for the increase in *de novo* NADP(H) synthesis observed in our experiments. In support of this idea, we observed that NADK levels are increased in the metastatic proficient 4T1 clone compared to the 4T07 clone that cannot effectively metastasize (Fig. 1g). Moreover, we observe the same trend in NADK levels when comparing human breast cancer cell lines derived from metastatic and non-metastatic sites (Fig. 1h). These observations raised the question of whether increased NADK levels could be generalizable to human cancer. Evaluation of human breast cancer samples showed an increase in NADK levels in metastases versus their matched primary tumors (Fig. 1i) indicating a direct relevance of *de novo* NADP(H) synthesis via NADK to metastasis formation in humans.

**Fig. 1.**
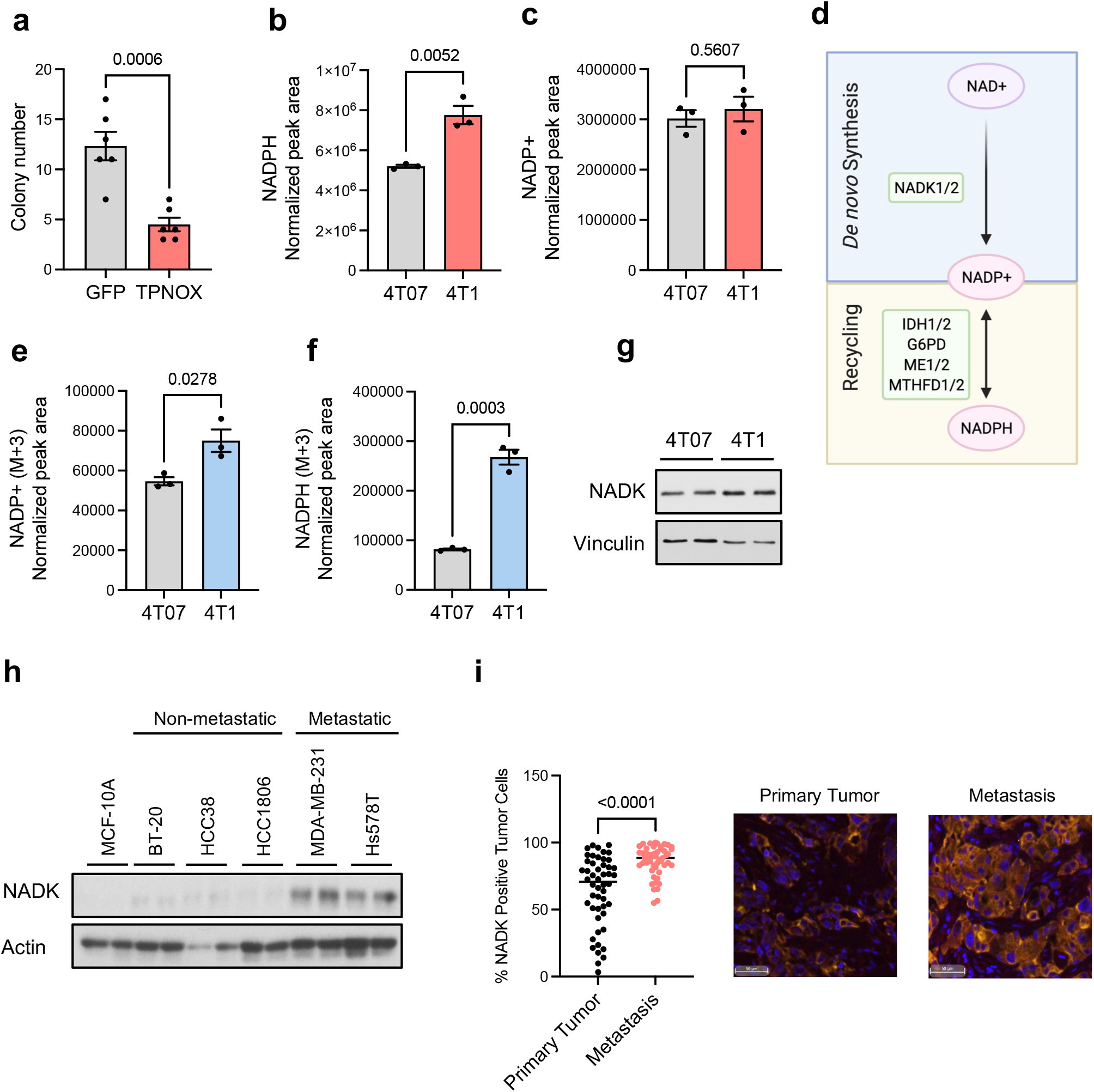
*De novo* NADP+ synthesis is upregulated in metastatic breast cancer cells. **a,** Soft agar colony formation assay in 4T1 cells expressing TPNOX or GFP (n=6). **b, c,** NADPH **(b)** and NADP+ **(c)** levels in 4T1 (broadly metastatic) and 4T07 (locally invasive) clones (n=3). **d,** Schematic representation of NADP(H) synthesis and recycling. **e, f,** *De novo* NADP+ **(e)** and NADPH **(f)** production from labeled NAM in 4T1 and 4T07 clones (n=3). **g,** NADK immunoblot in 4T1 and 4T07; representative image (n=4). **h,** NADK immunoblot in breast epithelial cells, non-metastatic breast cancer cells and metastatic breast cancer cells; representative image (n=4). **i,** Percentage of cells positive for NADK in human breast cancer primary tumor samples and matched metastasis (n=50) and representative images. All values are expressed as mean ± SEM.

Having shown that NADK is elevated in breast cancer metastases, we next probed its functional consequences. To test this, we silenced NADK in 4T1 cells as well as in a human cell line with high metastatic ability (MDA-MB-231 LM2 (21), referred to as LM2 hereafter). NADK silencing in both 4T1 and LM2 cells resulted in decreased total NADP(H) levels as well as *de novo* synthesized NADP+ in both cell lines (Extended Data Fig. 1a-c, Fig. 2a-d). Supporting the importance of *de novo* NADP+ synthesis for metastatic-like growth, we observed that NADK suppression in 4T1 or LM2 cells resulted in a pronounced decrease in their ability to grow in 3D, both in soft agar (Fig. 2e and 2f) and in the presence of an extracellular matrix hydrogel (basal-membrane extract – BME) (Fig. 2g and 2h). These data suggest that NADK suppression and the concomitant contraction of the total NADP(H) pools might impair metastasis formation. To directly evaluate how NADK might impact metastasis formation we performed lung colonization assays, which revealed that NADK suppression drastically reduced the ability of LM2 cells to effectively colonize the lung following a tail-vein injection (Fig. 2i).

**Fig. 2.**
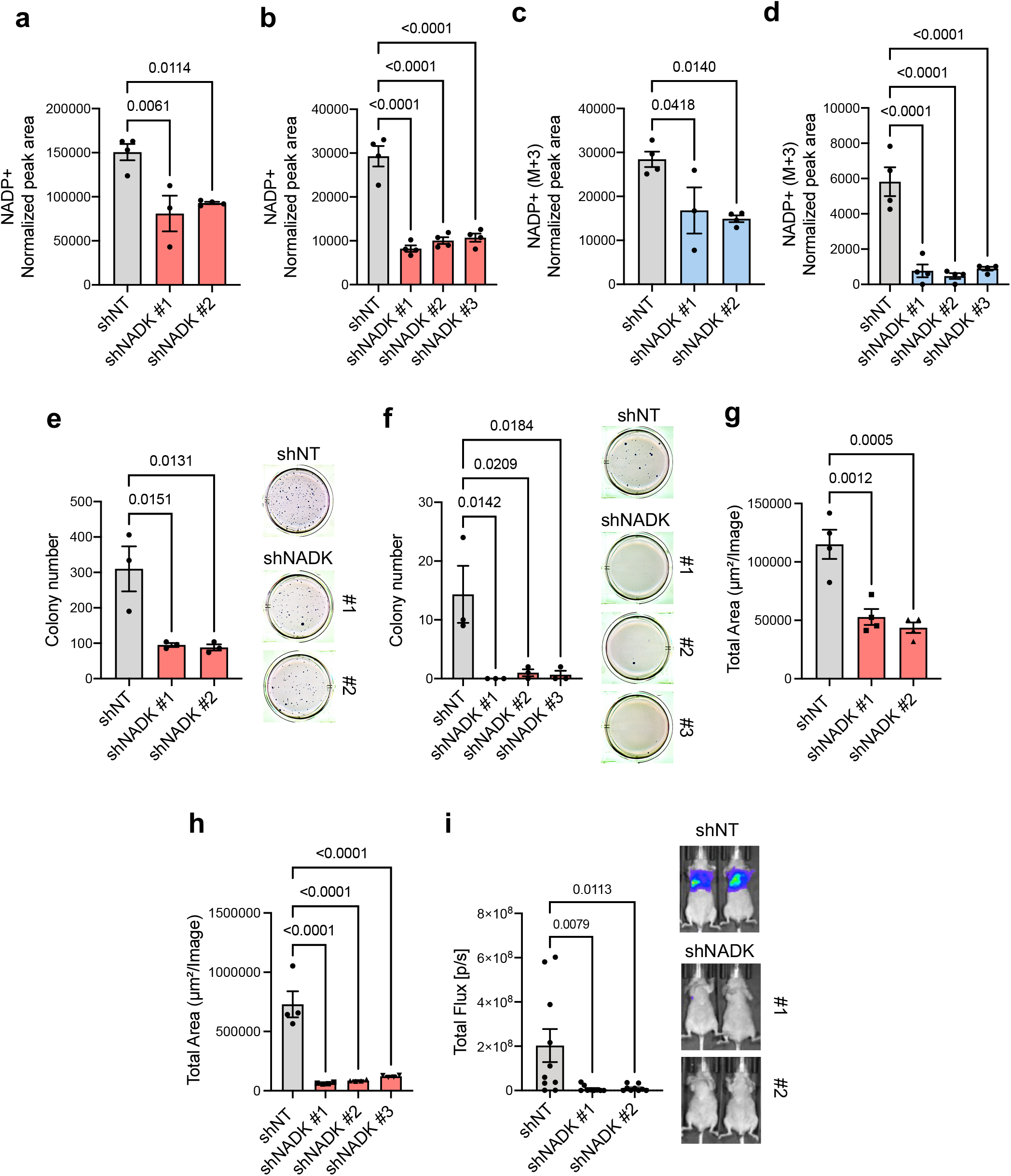
NADK mediates *de novo* NADP+ synthesis and enables metastatic colonization. **a, b,** NADP+ levels in 4T1 cells **(a)** and LM2 cells **(b)** with NADK knockdown for 3 days (n=3). **c, d,** *De novo* NADP+ production from labeled NAM in 4T1 cells **(c)** and LM2 cells **(d)** with NADK knockdown for 3 days (n=4). **e, f,** Quantification, and representative images, of soft agar colony formation assay in 4T1 **(e)** and LM2 cells **(f)** with NADK knockdown (n=3). **g, h,** Basal membrane extract 3D growth of 4T1 **(e)** and LM2 cells **(f)** with NADK knockdown (n=4). **i,** Lung colonization assay of LM2 cells with knockdown of NADK (n=10). All values are expressed as mean ± SEM.

To establish if the increase of NADK and consequent expansion of the NADP+ pools affect metastatic colonization by enabling ROS detoxification and/or fatty acid synthesis, we first measured ROS in 4T1 and LM2 cells upon NADK suppression. NADK knockdown in both cell lines resulted in suppression of antioxidant defences as shown by an expected decline in NADPH (Fig. 3a, Extended Data Fig. 2a) as well as a pronounced decline in reduced glutathione levels, which relies on the reducing power of NADPH to be recycled (Fig. 3b, Extended Data Fig. 2b). Accordingly, we observed a general increase in ROS (Fig. 3c, Extended Data Fig. 2c and 2d) and concomitant increase in the oxidation status of the cytosol as well as the mitochondria as evidenced by an increase in the oxidated fraction of peroxiredoxin 1 (PRDX1, cytosolic) and peroxiredoxin 3 (PRDX3, mitochondrial) upon NADK suppression (Fig. 3d). Moreover, in line with the importance of NADPH levels to power fatty acid synthesis we observed a decline in intracellular lipid levels upon NADK suppression (Fig. 3e and 3f). Importantly, NAD+ levels were not consistently altered upon NADK knockdown, excluding a potential effect of NADK in powering metastasis through regulation of NAD+ levels (Extended Data Fig. 1d and 1e). Having shown that NADK activity powers ROS detoxification and lipid synthesis in metastatic breast cancer cells, we sought to determine if deregulation of these pathways is at the root of the decline in metastatic ability upon NADK silencing. Supplementation with either lipids or a powerful antioxidant (N-acetylcysteine – NAC) alone were not sufficient to rescue the defect in BME growth elicited by the knockdown of NADK (Fig. 3g and Extended Data Fig. 2e). However, supplementation with both lipids and NAC gave both 4T1 and LM2 cells with NADK suppression a growth advantage both in 4T1 and LM2 cells (Fig. 3g and Extended Data Fig. 2e). Together, these observations suggest that expansion of the NADP(H) pools via *de novo* synthesis of NADP+ by NADK is an important feature for successful metastasis of breast cancer, by increasing redox power to fuel antioxidant defences and lipid synthesis.

**Fig. 3.**
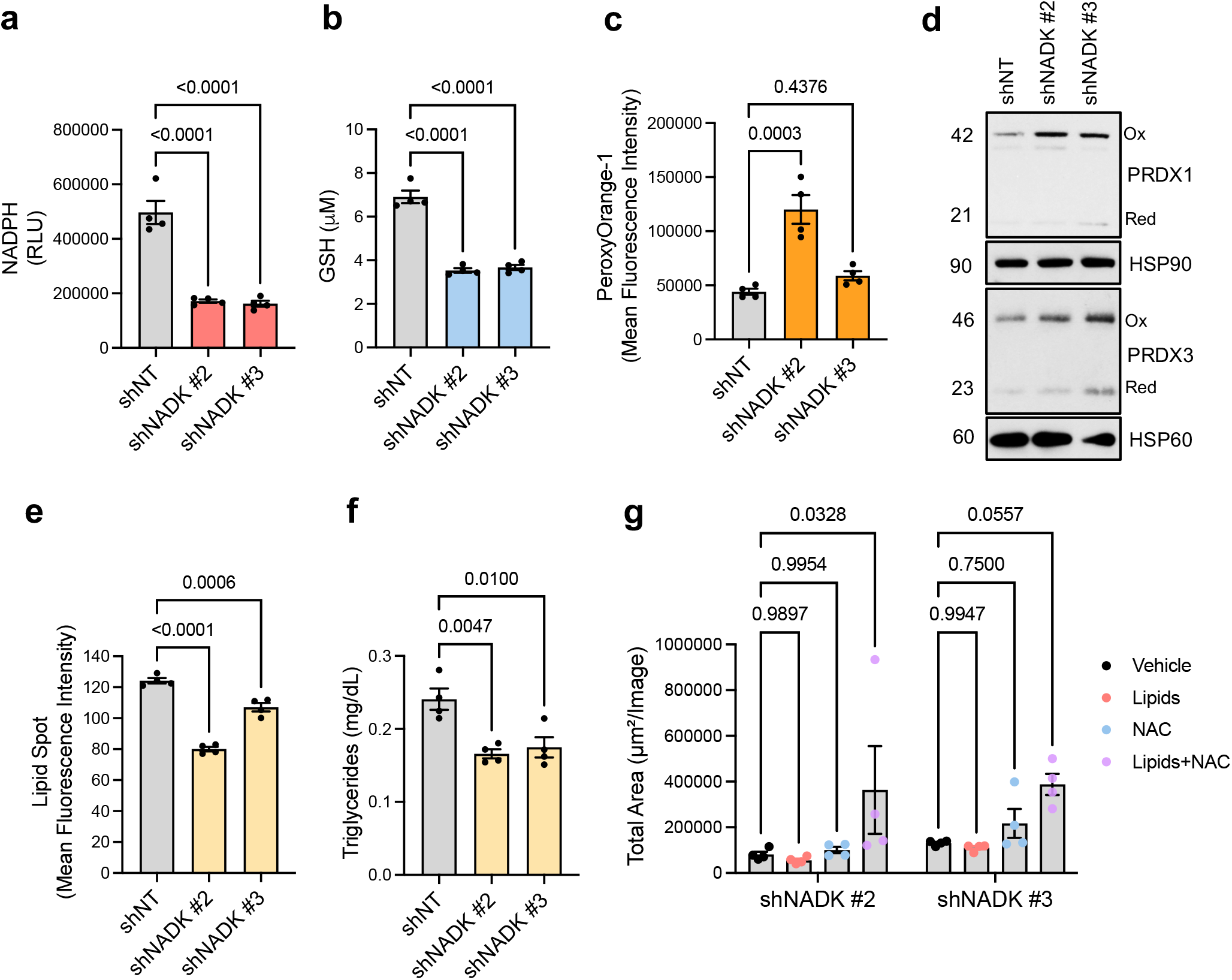
NADK increases NADP(H) pools to sustain redox and anabolic reactions and enable metastatic outgrowth. **a, b,** NADPH **(a)** and glutathione **(b)** levels in LM2 with NADK knockdown for 3 days (n=4). **c,** H_2_O_2_ levels in LM2 with NADK knockdown for 3 days (n=4). **d,** Redox status of PRDX1 and PRDX3 evaluated by immunoblot; representative image (n=4). **e, f,** Neutral lipid **(e)** and triglyceride levels **(f)** in LM2 with NADK knockdown for 3 days (n=4). **g,** Basal membrane extract 3D growth of LM2 cells with NADK knockdown supplemented with lipids or N-acetylcysteine (NAC) or the combination (n=4). All values are expressed as mean ± SEM.

Next, we sought to understand how NADK expression is regulated during tumour progression. Considering the importance of NADP(H) pools for redox control and anabolic processes, we reasoned that NADK induction might be an early event in the progression of breast cancers. To test this, we treated MCF10A, a breast epithelial cell line, with the well-established metastatic inducers TGFβ and TNFα (22). Consistent with their ability to promote EMT and the acquisition of metastatic traits (Fig. 4c), three days of TGFβ/TNFα treatment was sufficient to increase the levels of NADP+ in MCF10As (Fig. 4a). Moreover, TGFβ/TNFα treatment of HCC1806, a breast cancer cell line derived from a primary tumour, increased *de novo* production of NADP+ (Fig. 4b). Accordingly, we observed that NADK levels were upregulated in these models as well as in HCC38, another breast cancer cell line derived from a primary tumour, upon treatment with TGFβ/TNFα (Fig. 4c). We had previously demonstrated that in order for breast cancer cells to acquire metastatic-like properties in response to TGFβ/TNFα, an epigenetic reprogramming regulated by histone H3 chaperones and the non-canonical H3.3 variant needs to occur (23). Analysis of publicly available H3.3 ChIP-seq in LM2 cells (23) demonstrated an enrichment of H3.3 at the promoter region of NADK (Extended Data Fig. 3a), suggesting that NADK regulation by metastatic signals such as TGFβ and TNFα might be mediated by H3.3 deposition. In support of this hypothesis, ChIP-PCR analysis demonstrated H3.3 incorporation into the *NADK* promoter was elevated in response to TGFβ/TNFα treatment (Fig. 4d). H3.3 is thought to gap fill the DNA in response to stimuli that decrease the incorporation of canonical histones into chromatin (24). Suppression of CAF-1, the histone chaperone complex in charge of depositing canonical histones into chromatin, has previously been shown to be sufficient to trigger this H3.3 gap filling mechanism in breast cancer (23). Thus, we reasoned that if H3.3 gap filling is mediating the induction in NADK in response to TGFβ and TNFα, suppression of the CAF-1 complex alone should induce the increase in NADK. Suppressing the CAF-1 complex via shRNA-mediated silencing of its subunit CHAF1B resulted in a similar increase in H3.3 enrichment at the promoter of NADK, increased levels of NADK and concomitantly increased NADP(H) levels in non-metastatic breast cancer cells (Extended Data Fig. 3b-f), mirroring the effects of TGFβ/TNFα treatments. Moreover, suppression of HIRA, the chaperone responsible for incorporation of H3.3 onto chromatin in genic regions, ablated the ability of TGFβ and TNFα to induce NADK levels and to increase NADP(H) levels (Fig. 4e-g, Extended Data Fig. 4a and 4b). Finally, because NADK levels are elevated in metastatic breast cancer cells (Fig. 1h) and H3.3 was enriched at the NADK promoter of LM2 cells (Extended Data Fig. 3a), we evaluated if suppressing HIRA in this context would also affect NADK levels and the NADP(H) pools. Our data show that suppression of HIRA in breast cancer cells with metastatic ability decreased H3.3 enrichment at the NADK promoter, resulting in a decrease in NADK mRNA and protein levels and ultimately leads to the contraction of NADP(H) pools in these cells (Fig. 4h-l, Extended Data Fig. 4c-e). These data show that metastatic signals drive NADK expression through deposition of H3.3 into the NADK promoter resulting in its activation, illustrating a complete epigenetic mechanism that regulates the NADP(H) pools.

**Fig. 4.**
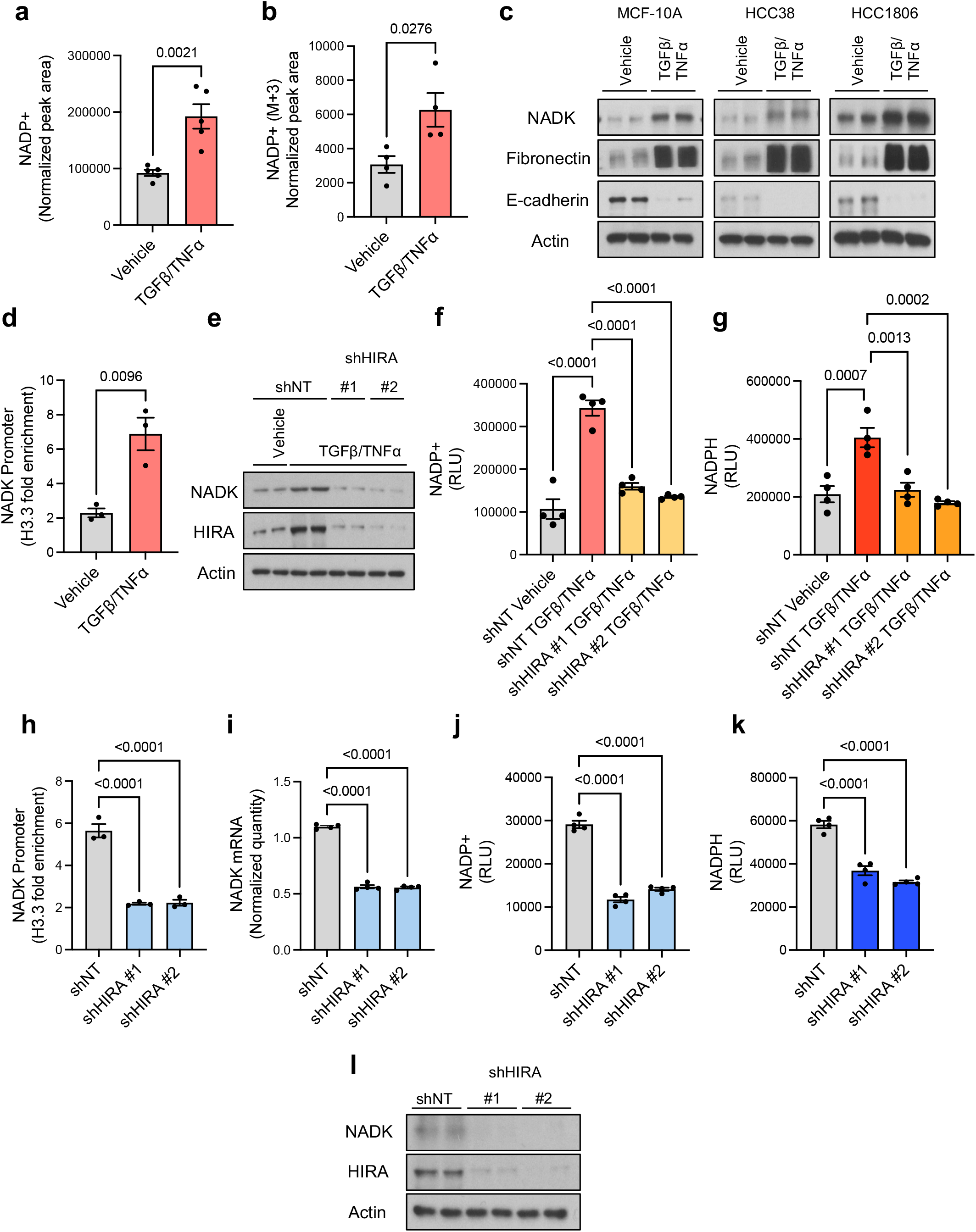
H3.3 incorporation into chromatin in response to metastatic signaling drives NADK expression in breast cancer. **a,** NADP+ total levels in MCF-10A cells treated with TGFβ + TNFα for 3 days (n=5). **b,** *De novo* NADP+ production from labeled NAM in HCC1806 cells treated with TGFβ + TNFα for 3 days (n=4). **c,** NADK immunoblot in MCF-10A, HCC38 and HCC1806 cells treated with TGFβ + TNFα for 5 days; representative images (n=4). **d,** H3.3 enrichment at the *NADK* promoter in MCF-10A treated with TGFβ + TNFα for 3 days; fold enrichment was determined using immunoglobulin G (IgG) as a control for the ChIP (n=3). **e,** NADK immunoblot in HCC1806 cells with HIRA knockdown treated with TGFβ + TNFα for 5 days; representative image (n=4). **f, g,** NADP+ **(f)** and NADPH **(g)** levels in HCC1806 cells with HIRA knockdown treated with TGFβ + TNFα for 5 days (n=4). **h,** H3.3 enrichment at the *NADK* promoter in LM2 cells with HIRA knockdown for 3 days; fold enrichment was determined using immunoglobulin G (IgG) as a control for the ChIP (n=3). **i,** NADK mRNA quantity in LM2 cells with HIRA knockdown for 3 days (n=4). **j, k,** NADP+ **(j)** and NADPH **(k)** levels in LM2 cells with HIRA knockdown for 3 days (n=4). **l,** NADK immunoblot in LM2 cells with HIRA knockdown for 3 days; representative image (n=4). All values are expressed as mean ± SEM.

Taken together, our results show that NADK upregulation and the consequent expansion in the NADP(H) pools play a central role in the ability of breast cancer cells to colonize distal organs and thrive as metastatic lesions by powering anabolic reactions and antioxidant systems. Mechanistically, we demonstrated that this occurs in response to metastatic signals, such as TGFβ and TNFα, via chromatin remodelling that impinges on the histone H3.3 variant. Although more in-depth studies are necessary to fully determine the scope of H3.3-mediated regulation of metabolic pathways in breast cancer metastasis, this study adds NADK and *de novo* NADP(H) synthesis to the growing list of essential metabolic adaptations that enable metastases to thrive. Our results suggest that NADK might be an important new and effective therapeutic target for advanced breast cancers and metastatic disease.

## Contributions

A.P.G conceived the project. T.S. and D.I. performed all the molecular biology experiments, the 3D growth-related experiments, the NADP(H) and GSH measurements, the mechanistic experiments, prepared cells for mass spectrometry and isotope tracing analysis and assisted on all other experiments. M.R.M. performed the mass spectrometry and isotope tracing analysis. N.P.W. performed the ROS measurements and the redox blots. S.D. performed the lipid analysis. V.L. performed the mouse experiment. E.A. performed the NADK analysis in human breast cancer patient samples. A.P.G. performed the ChIP experiments. A.P.G., M.R.M, G.M.D. and E.L-L. supervised the project. A.P.G., D.I., T.S., N.P.W., S.D., V.L., E.A. and M.R.M analysed the data. The manuscript was written by A.P.G. and edited by D.I., T.S., N.P.W., G.M.D and M.R.M. All authors discussed the results and approved the manuscript.

## Acknowledgements

We are grateful to members of the Gomes and DeNicola Laboratories as well as Dr. Gerta Hoxhaj for critical input on this project. We are also thankful to Dr. William Schiemann for the 4T1 clones and Dr. Joan Massague for the MDA-MB-231 LM2 cells. The Gomes Lab is supported by a Pathway to Independence Award to A.P.G. from NCI (R00CA218686), a New Innovator Award from OD/NIH (DP2AG0776980), the American Lung Association, the Florida Health Department Bankhead-Coley Research Program, the Florida Breast Cancer Foundation, and the George Edgecomb Society of Moffitt Cancer Center. T.S. was supported by the NIH F31 pre-doctoral fellowship F31CA220750. M.R.M is supported by HHMI Hanna H. Gray Fellows Program Faculty Phase (Grant #GT15655). G.M.D is supported by the NIH/NCI (R37CA230042).This work was also supported by Moffitt’s Cancer Center Support Grant from NCI (P30CA076292) through the Analytic Microscopy Core. No potential conflicts of interest were disclosed by the authors.

## Methods

Email contact for reagent and resource sharing

### Cell Lines

HCC38, HCC1806, BT20, MDA-MB-231, Hs578t and MCF10A cells were obtained from American Type Culture Collection (ATCC). 4T1 and 4T07 mouse breast cancer cell line clones were originally derived by Dr. F. Miller^19^ and obtained from Dr. William Schiemann. MDA-MB-231 LM2 (referred to as LM2) subclone^18^ was obtained from Dr. Massague’s lab. HEK293T cells were obtained from GenHunter. All cell line were maintained at 37°C and 5% CO_2_ in the presence of 100 unit/ml penicillin and 100 μg/ml streptomycin (Gibco) and were routinely tested for mycoplasma using MycoAlert mycoplasma detection kit (Lonza). HCC38, HCC1806, BT20, MDA-MB-231, 4T1 and 4T07 cells were cultured in RPMI-1640 medium supplemented with 10% FBS. Hs578T cells were maintained in high glucose DMEM with 0.01 mg/ml insulin and 10% FBS, while LM2 and HEK293T cells were cultured in high glucose DMEM with 10% FBS. MCF10A cells were maintained in DMEM:F12 media supplemented with 5% horse serum (Gibco), 10 μg/mL insulin (Sigma-Aldrich), 100 ng/mL cholera toxin (Sigma-Aldrich), 20 ng/mL EGF (Peprotech), and 0.5 mg/mL hydrocortisone (Sigma-Aldrich). The cells used for the experiments in this manuscript always tested negative for mycoplasma.

### Mice

Female nu/nu athymic mice, 4 to 6 weeks old, were obtained from Envigo. The animals were allowed acclimate for at least 7 days before experiments. The mice were maintained at Weill Cornell Medicine vivarium under standard husbandary conditions with unrestricted access to food and water in compliance with the Weill Cornell Medicine Institutional Animal Care and Use Committee (IACUC) protocols. The animal room was maintained with a 12 hours light-dark cycle at 21-23 °C, around 50% humidity and. The standard chow, PicoLab Rodent Diet 5053 (Labdiet, Purina) containing 20% protein and 5% fat, was used. The maximum tumor size allowed—20 mm or 2.5 cm^3^, or 10% of the animal’s body weight—by the IACUC protocol was not exceeded.

## METHOD DETAILS

### Cell Culture Treatments

Some cells were treated with 5 ng/ml of recombinant human TGFβ (PeproTech) and 5 ng/ml of recombinant human TNFα (PeproTech) as metastatic inducers for the indicated time periods. When cells with genetic suppression of a gene of interest were also treated with TGFβ1 and TNFα, the cells were transduced and selected as described below and then treated with 5 ng/ml of recombinant human TGFβ (PeproTech) and 5 ng/ml of recombinant human TNFα (PeproTech).

### 3D Soft Agar Colony Formation Assay

As the bottom layer a coating of 1:1 ratio of 2 × DMEM (Millipore): 1.2% SeaPlaque Agarose (Lonza) was applied to 6-well plates, and allowed to set overnight. The top layer was applied the next day (1:1 ratio of 2 × DMEM and 0.7% agarose) containing 5,000 of the indicated cells per well. The next day 0.5 mL 1 × DMEM with selection antibiotic was added to each well. The media was changed once a week and the colonies were allowed to grow for 4-6 weeks. To visualize and quantify 3D colonies, cells were stained in 0.005% Crystal violet/10% Ethanol aqueous solution for 3 hours, and de-stained in water overnight. Colonies were counted by eye.

### Basal Membrane Extract (BME) 3D growth assay

96-well plates were coated with 40 μL of growth-factor reduced BME (Cultrex), and plates were allowed to set for at least an hour at 37 °C. 2,000 cells/well in 100 μL media supplemented with 2% FBS, 2% BME and appropriate selection antibiotics was layered on top of the BME coating. The plates were imaged with Incucyte Live Imaging system every 12 hours. The cells were fed fresh media supplemented with 2% FBS every 4 days. Where indicated cell media was also supplemented with 2 mM N-acetyl-L-cysteine (NAC, Sigma-Aldrich), or lipids (12 μg/ml low density lipoprotein (Kalen Biomedical) and 50 μM oleate (Sigma-Aldrich)) or in combination. Colony growth was analyzed using Incucyte software (v. 2021A) as total area covered by the colonies.

### Isotope Tracing of *de novo* NADP+ Generation

Cells were grown in nicotinamide (NAM) free media with 10% dialyzed serum (Sigma-Aldrich) to 60-80% confluency in 6 cm plates. Prior to harvesting cells were treated with 32 μM [2H]NAM for 2 hours. The media was aspirated and 1 mL of ice-cold 40% MeOH/40% acetonitrile/20% water/0.5% formic acid mixture was added to the plates and incubated for 30 seconds before 80 μL of 2M NH_4_HCO_3_ was added to neutralize. Plates were moved to −20 °C for 30 min, before cells were scraped off and moved into microcentrifuge tubes. The tubes were incubated on dry ice for 5 min centrifuged at 16,000 g for 10 minutes at 4 □C. Supernatants were collected and frozen at −80 □C until further processing. Within 24 hours before analysis by liquid chromatography coupled to a mass spectrometer (LC-MS), the supernatants were centrifuged again at 16,000 g for 20 minutes to remove any debris. The LC–MS method was based on hydrophilic interaction chromatography (HILIC) coupled to the Q Exactive PLUS mass spectrometer (Thermo Scientific). The LC separation was performed on a XBridge BEH Amide column (150 mm 3 2.1 mm, 2.5 mm particle size, Waters, Milford, MA). Solvent A is 95%: 5% H2O: acetonitrile with 20 mM ammonium bicarbonate, and solvent B is acetonitrile. The gradient was 0 min, 85% B; 2 min, 85% B; 3 min, 80% B; 5 min, 80% B; 6 min, 75% B; 7 min, 75% B; 8 min, 70% B; 9 min, 70% B; 10 min, 50% B; 12 min, 50% B; 13 min, 25% B; 16 min, 25% B; 18 min, 0% B; 23 min, 0% B; 24 min, 85% B; 30 min, 85% B. The following parameters were maintained during the LC analysis: flow rate 150 mL/min, column temperature 25 °C, injection volume 10 μL and autosampler temperature was 5 °C. For the detection of metabolites, the mass spectrometer was operated in both negative and positive ion mode. The following parameters were maintained during the MS analysis: resolution of 140,000 at m/z 200, automatic gain control (AGC) target at 3e6, maximum injection time of 30 ms and scan range of m/z 75-1000. If variation in cell numbers was observed between conditions, the data were normalized to the protein content. The MAVEN software was used to analyze the data, and natural 13C abundance was taken into account to correct all labeling patterns using AccuCor^1^. When normalization was required due to variation in cell number in between samples, the original data were normalized to protein content.

### NAD+ and NADP(H) measurement

Whenever data are presented as normalized peak area, cells grown normally without any tracers were harvested and analyzed through LC-MS as described above in the isotope tracing section to determine the NAD+ or NADP(H) levels in the cells. Whenever data are presented as relative light unit (RLU), NADP/NADPH-Glo™ Assay (Promega), a luminescence-based method, was used to measure NADP+ and NADPH of 100,000 cells on 96-well assays according to manufacturer’s instructions. The luminescence was measured using a Varioskan microplate reader (Thermo Scientific).

### Generation of Stable Cell Lines

For gene silencing experiments, shNT (shGFP—TRCN0000072181), shNADK#1 (TRCN0000037700), shNADK#2 (TRCN0000199808), shNADK#3 (TRCN0000199040), shNadk#1 (TRCN0000297518), shNadk#2 (TRCN0000278616), shCHAF1B #1 (TRCN0000074279), shCHAF1B #2 (TRCN0000074278) were obtained from Sigma, and shHIRA #1 (TRCN0000020514) and shHIRA #2 (TRCN0000020515) were obtained from OpenBiosystems. pMD2.G (Addgene plasmid 12259) and psPAX2 (Addgene plasmid 12260) constructs were co-transfected with each construct in HEK293T cells using X-tremeGENE HP (Roche) according to manufacturer’s protocols to produced viral particles expressing the shRNA of interest. The media was refreshed 24 hours after transfection and media containing virus was collected 72 hours after transfection. Filtered virus was used for transductions in the presence of 8 μg/mL polybrene (Sigma-Aldrich). Cells were selected with 2 μg/mL puromycin (Sigma-Aldrich) starting 24 hours after transduction, and 2 μg/mL puromycin (Sigma-Aldrich) was maintained in their growth media for the duration of the experiments.

TPNOX and the control GFP cDNAs were custom made and purchased from VectorBuilder (C-terminally FLAG-tagged lentiviral constructs under EF1a promoter)., pRSV-Rev (Addgene plasmid 12253), pMDLg/pRRE (Addgene plasmid 12251) and pMD2.G (Addgene plasmid 12259) co-transfected with each construct in HEK293T cells using X-tremeGENE HP (Roche) according to manufacturer’s protocols to produced viral particles expressing the cDNAs of interest. The media was refreshed 24 hours after transfection and media containing virus was collected 72 hours after transfection. Filtered virus was used for transduction of 4T1 cells in the presence of 8 μg/mL polybrene (Sigma-Aldrich). 4T1 cells were selected with 5 μg/ml of Blasticidin⋅HCl (Thermo Fisher) starting 24 hours after transduction, and 5 μg/ml of Blasticidin⋅HCl (Thermo Fisher) was maintained in their growth media for the duration of the experiments.

### Immunoblots for Total Cell Lysates

Proteins were harvested with 10% TCA solution (10% trichloroacetic acid, 25 mM NH_4_OAc, 1 mM EDTA, 10 mM Tris⋅HCl pH 8.0), and pelleted proteins were resolubilized in a 0.1 M Tris⋅HCl pH 11 solution containing 3% SDS by boiling for 10-15 minutes. Protein concentration was determined with DC Protein Assay kit II (BioRad)and 20 μg total protein per sample was run on SDS-PAGE under reducing conditions, and then transferred from the gels to nitrocellulose membranes (GE Healthcare) electrophoretically. The membranes were blocked in 5% milk and then incubated with the primary antibodies overnight at 4°C. The antibodies used to detect the proteins of interest were: E-Cadherin (610181 - BD Biosciences, Dilution 1:1000), Fibronectin (ab2413 – Abcam, Dilution 1:5000), NADK (55948S - Cell Signaling, Dilution 1:500), CHAF1B (HPA021679 – Sigma-Aldrich, Dilution 1:1000), HIRA (ab129169 – Abcam, Dilution 1:500), vinculin (V9264 – Sigma-Aldrich, Dilution 1:10,000) and Actin (sc1615 - Santa Cruz, Dilution 1:10,000). The membranes were then incubated with the appropriate horseradish peroxidase– conjugated (HRP) anti-rabbit (NA934-Cytiva, Dilution 1:10,000), anti-mouse (NA931-Cytiva, Dilution 1:10,000), or anti-goat (AP180P-Millipore, Dilution 1:10,000) immunoglobulin for 2 hours at room temperature. Amersham ECL detection system (GE Healthcare) was utilized to develop the signals.

### Redox Immunoblotting

PRDX1 and PRDX3 oxidation states were assessed in accordance with a previously established protocol^2^. For each biological replicate, 500,000 cells were seeded overnight on a 6-well plate. Cells were then washed twice with cold PBS (Hyclone) and overlaid with 200 μL of alkylation buffer (40 mM HEPES [VWR], 50 mM NaCl [Fisher Scientific], 1 mM EGTA [VWR], Pierce Protease Inhibitor [Fisher Scientific]) supplemented with 200 mM N-ethylmaleimide (NEM; Alfa Aesar). Cells were incubated for 10 minutes at room temperature and then 20 μL of 10% CHAPS detergent was added and cells incubated at room temperature for an additional 10 minutes to lyse cells. Lysates were then collected, vortexed, and cleared by centrifugation for 15 minutes at 17,000 g at 4 °C. Supernatants were isolated to quantify protein and 5-10 μg protein samples were mixed with a 4X non-reducing buffer prior to separation by SDS-PAGE. Separated proteins were transferred to a nitrocellulose membrane and incubated with antibodies recognizing HSP60 (Cell Signaling Technologies, D307), HSP90 (Cell Signaling Technologies, 4874S), PRDX1 (Cell Signaling Technologies, D5G12), or PRDX3 (Abcam, ab73349). HRP-conjugated secondary antibodies and enhanced chemiluminescence were used for redox immunoblotting.

### NADK fluorescent immunohistochemistry (IHC)

Immunostaining for NADK was performed on paraffin-embedded FFPE tumor tissue sections (TMA slide, Serial #BR10010f; US BioMax). The slide was melted at 70 °C for 30 min and was further de-paraffinized using xylene and rehydrated in serial alcohol washes. The slide was pressure cooked at 15 PSI for 15 min in a 1X DAKO antigen retrieval buffer (Agilent Technologies). Tissue sections were subject to two 5-min standing washes in PBS prior to blocking in 1X Carb-Free Blocking Solution (Vector Labs) for 3h at room temperature. The slide was next washed twice and incubated with anti-NADK primary antibody (15548-1-AP dilution:1:50; Proteintech) overnight at 40 °C. Next day, the slide was washed with PBS three times and incubated with AlexaFluor 647 (Cy5) anti-rabbit secondary antibody (A32733 dilution:1/250; Invitrogen) and eFluor 570 (Cy3) Anti-Pan Cytokeratin (41-9003-82 dilution:1/400; Invitrogen) in dark for 3 hours at room temperature. The slide was next washed and mounted with Vectashield + DAPI (Vector Laboratories). TMA was scanned on the Aperio FL (Leica Biosystems) using DAPI, Cy3 and Cy5 fluorescent filters and the whole slide FL image scan was loaded into HALO Image Analysis Platform v3.3 (Indica Labs, Albuquerque, NM) for quantitative image analysis. The TMA was segmented in HALO and each core is reviewed for lifted and blurred areas of tissue and an inclusion or exclusion for analysis annotations are created. A random forest machine learning classifier was trained on multiple tissue cores using the PCK and DAPI channels to identify tumor vs non-tumor tissue. The tissue is then segmented into individual cells using the DAPI marker which stains cell nuclei. For each marker, a positivity threshold within the nucleus or cytoplasm are determined per marker based on visual FL staining. After setting a positive fluorescent threshold for each staining marker, the slide is analyzed with the algorithm. The generated data includes positive cell counts for each fluorescent marker and percent of cells positive for the marker. Along with the summary output, a per-cell analysis can be exported to provide the marker status and fluorescent intensities of every individual cell within an image.

### Gene Expression Analysis

Total RNA was extracted from cells grown on 6 cm dishes until 70-80% confluency using the PureLink RNA isolation kit (Life Technologies). To digest contaminating DNA, isolated RNA was treated with DNAse I (Amplification grade, Sigma-Aldrich). cDNA was synthesized with iSCRIPT cDNA synthesis kit (BioRad) and quantitative PCR (qPCR) using SYBR green master mix (Life Technologies) was performed on a QuantStudio6 Real-Time PCR system (Life Technologies, software version v1.3). Tata Binding Protein (TBP) expression were used to normalize NADK expression levels. The primer sequences are:

TBP_forward: GAGCCAAGAGTGAAGAACAGTC

TBP_reverse: GCTCCCCACCATATTCTGAATCT

NADK_forward: GAAGCAAGGAACACAGCATG

NADK_reverse: CTCTCAAACCAGTCGCTCAC

### Lung Colonization in Mice

Female nu/nu athymic mice were injected intravenously (tail-vein) with 100,000 LM2 cells with knockdown of NADK. For each experimental group 10 mice were used. IVIS Spectrum CT Pre-Clinical In Vivo Imaging System (Perkin-Elmer) was used to monitor the metastases to evaluate lung colonization. The luminescence was quantified 6 weeks after the injections using the Living Image Software (v4.5, Perkin-Elmer). The Weill Cornell Medicine IACUC guidelines were followed during the experiments.

### Flow Cytometry Analyses of ROS

For all analyses of ROS, 500,000 cells of each biological replicate were seeded overnight on a 6-well plate. Total cellular ROS levels were determined with the fluorescent dye, CellRox Deep Red (Invitrogen), according to the manufacturer’s protocol. Briefly, cells were incubated in 1mL of fresh media for 4 hours, at which point 2 μL of 2.5mM CellROX Deep Red was added to each well for a final concentration of 5 μM. Cells were incubated with CellROX Deep Red for 30 minutes and then collected for analysis. Cellular H_2_O_2_ levels were determined using the fluorescent dye, Peroxy Orange-1 (Fisher Scientific). Cells were incubated in 1 mL of fresh media for 4 hours, then washed twice with PBS (Hyclone) and incubated in 1 mL of 5 μM Peroxy Orange-1 for 40 minutes. The fluorescence of dye-loaded cells was determined by flow cytometry with a BD Accuri C6 Plus Flow Cytometer (BD Biosciences). An allophycocyanin (APC) channel was used for analyses of CellROX Deep Red fluorescence, whereas a phycoerythrin (PE) channel was used for analyses of Peroxy Orange-1 fluorescence. The mean fluorescence intensity of 10,000 discrete events were calculated for each biological replicate.

### GSH measurement

Cells were seeded and grown to 80% confluency on 96-well plates. Luminescence based GSH-Glo™ Assay (Promega) was used to determine the cellular glutathione (GSH) levels of 100,000 cells on 96-well according to the manufacturer’s manual. The luminescence was measured using a Varioskan microplate reader (Thermo Scientific).

### Lipid Droplet Stain

500,000 cells were seeded on 6-well plates overnight. Cells were incubated with LipidSpot™ 610 (Biotium) for 30 minutes at 37 °C to stain lipid droplets according to manufacturer’s instructions. Samples were analyzed by SONY SH800S flow cytometer (SONY). The fluorescence of LipidSpot™ 610 was determined using Texas Red channel (excitation/emission at ~592/638 nm). Acquired FCS files were exported and analyzed using FlowJo software (v10.0.7;BD). Cell aggregates and debris were excluded from the analysis based on a dual-parameter dot plot in which the pulse ratio (signal height/y-axis vs. signal area/x-axis) was displayed. The median fluorescence intensity of 20,000 events were calculated for each biological replicate.

### Triglyceride Measurement

Cells were seeded on 6 cm dishes and grown to 80% confluency. Triglyceride Colorimetric Assay Kit (Cayman Chemical) was used to determine the triglyceride levels in cells according to the manufacturer’s manual. The absorbance was measured using a Varioskan microplate reader (Thermo Scientific)

### H3.3 Signal Tracks

ChIP-seq data for H3.3 in LM2 cells^3^ (GEO: GSE120313) was used to visualize H3.3 signal tracks for the NADK gene using Integrated Genome Browser (IGB)^4^.

### ChIP-PCR

ChIP-IT Express (Active Motif) kit was used to perform chromatin immunoprecipitation (ChIP) according to the manufacturer’s instructions. The chromatin was sheared enzymatically for 10 minutes (ChIP-IT Express Enzymatic Shearing Kit, Active Motif). Instead of the magnetic beads from the kit, Dynabeads (1:1 protein A to protein G; Life Technologies) were used. Anti-H3.3 (17-10245, Millipore) or anti-Rabbit IgG (sc-2027, Santa Cruz) antibodies were used for the immunoprecipitation. Chromatin IP DNA purification kit (Active Motif) was used to purify DNA and qPCR was performed with SYBR Green master mix (Life Technologies) on QuantStudio Pro 6 (Life Technologies) using the primer set for NADK listed below, corresponding to the DNA sequence on NADK promoter indicated with a red arrow on Extended Data Figure S3a:

NADK-ChIP-forward: CGCAGTTCCAACAAACACTAC

NADK-ChIP-reverse: GAGGACTCGGGGAGTTGG

### Statistical Analysis

GraphPad Prism 7 or 9, and Microsoft Excel 2013 or 365 were utilized for data analyses. was used to determine significance When two conditions were compared two-tailed Student’s t test, and for experiments with more than two conditions ANOVA analyses were used to determine significance. Data are from at least three independent experiments and represented as the mean ± SEM (standard error of the mean) of individual data points. Number of replicates and animals are reported in the figure legends. Normal distribution of samples was not determined, but similar variances between groups were observed in all experiments.

## Data Availability

The raw data supporting each figure and the raw images for the western blots can be found in the corresponding Source Data files. The accession number for the raw ChIP-sequencing data that is previously published ^3^ and publicly available is GEO: GSE120313. This data has been used to generate H3.3 tracks for the NADK gene.

## Code Availability

No code was created for this manuscript.

## Extended Data Figure Legends

**Extended Data Fig. 1 – NADK regulates NADP(H) levels without altering NAD+ levels in metastatic breast cancer cells. a, b,** NADK immunoblot in 4T1 **(a)** and LM2 cells with NADK knockdown for 3 days; representative image (n=4). **c,** NADPH levels in 4T1 cells with NADK knockdown for 3 days (n=3). **d, e,** NAD+ levels in 4T1 cells **(d)** and LM2 cells **(e)** with NADK knockdown for 3 days (n=3). All values are expressed as mean ± SEM.

**Extended Data Fig. 2 – NADK is an important regulator of redox homeostasis in murine breast cancer metastatic cells. a, b,** NADPH **(a)** and glutathione **(b)** levels in 4T1 cell with NADK knockdown for 3 days (n=4). **c, d,** ROS levels measured by PeroxyOrange-1 **(c)** and CellROX Deep Red **(d)** in 4T1 cell with NADK knockdown for 3 days (n=4). **e,** Basal membrane extract 3D growth of 4T1 cells with NADK knockdown supplemented with lipids or N-acetylcysteine (NAC) or the combination (n=4). All values are expressed as mean ± SEM.

**Extended Data Fig. 3 – Suppression of the CAF-1 complex triggers H3.3 deposition at the *NADK* promoter to expand NADP(H) pools. a,** H3.3 signal track of H3.3 ChIP-seq analysis in LM2 cells. **b,** H3.3 enrichment at the *NADK* promoter in MCF-10A with CHAF1B suppression for 3 days; fold enrichment was determined using immunoglobulin G (IgG) as a control for the ChIP (n=3). **c, d,** NADK immunoblots in HCC38 **(c)** and HCC1806 **(d)** with CHAF1B suppression for 3 days; representative images (n=4). **e, f,** NADP+ and NADPH levels in HCC38 **(e)** and HCC1806 cells **(f)** with CHAF1B knockdown for 3 days (n=4). All values are expressed as mean ± SEM.

**Extended Data Fig. 4 – HIRA-mediated H3.3 deposition regulates NADK levels and NADP(H) total pools. a,** NADK immunoblot of HCC38 cells with HIRA knockdown treated with TGFβ + TNFα for 5 days; representative image (n=4). **b,** NADP+ and NADPH levels in HCC38 cells with HIRA knockdown treated with TGFβ + TNFα for 5 days (n=4). **c,** NADK mRNA quantity in Hs578T cells with HIRA knockdown for 3 days (n=4). **d,** NADK immunoblot in Hs578Tcells with HIRA knockdown for 3 days; representative image (n=4). **e,** NADP+ and NADPH levels in Hs578T cells with HIRA knockdown for 3 days (n=4). All values are expressed as mean ± SEM.

## References

1. Valastyan S, Weinberg RA. Tumor metastasis: molecular insights and evolving paradigms. Cell. 2011;147(2):275–92. doi: 10.1016/j.cell.2011.09.024. PubMed PMID: 22000009; PMCID: PMC3261217.

2. Chiang AC, Massague J. Molecular basis of metastasis. N Engl J Med. 2008;359(26):2814–23. Epub 2008/12/26. doi: 10.1056/NEJMra0805239. PubMed PMID: 19109576; PMCID: PMC4189180.

3. Gupta GP, Massague J. Cancer metastasis: building a framework. Cell. 2006;127(4):679–95. Epub 2006/11/18. doi: 10.1016/j.cell.2006.11.001. PubMed PMID: 17110329.

4. Drapela S, Gomes AP. Metabolic requirements of the metastatic cascade. Curr Opin Syst Biol. 2021;28. Epub 2021/10/26. doi: 10.1016/j.coisb.2021.100381. PubMed PMID: 34693082; PMCID: PMC8535854.

5. Bergers G, Fendt SM. The metabolism of cancer cells during metastasis. Nat Rev Cancer. 2021;21(3):162–80. Epub 2021/01/20. doi: 10.1038/s41568-020-00320-2. PubMed PMID: 33462499; PMCID: PMC8733955.

6. Broadfield LA, Pane AA, Talebi A, Swinnen JV, Fendt SM. Lipid metabolism in cancer: New perspectives and emerging mechanisms. Dev Cell. 2021;56(10):1363–93. Epub 2021/05/05. doi: 10.1016/j.devcel.2021.04.013. PubMed PMID: 33945792.

7. Gill JG, Piskounova E, Morrison SJ. Cancer, Oxidative Stress, and Metastasis. Cold Spring Harb Symp Quant Biol. 2016;81:163–75. Epub 2017/01/14. doi: 10.1101/sqb.2016.81.030791. PubMed PMID: 28082378.

8. Tasdogan A, Ubellacker JM, Morrison SJ. Redox Regulation in Cancer Cells during Metastasis. Cancer Discov. 2021;11(11):2682–92. Epub 2021/10/16. doi: 10.1158/2159-8290.CD-21-0558. PubMed PMID: 34649956; PMCID: PMC8563381.

9. Martin-Perez M, Urdiroz-Urricelqui U, Bigas C, Benitah SA. Lipid metabolism in metastasis and therapy. Current Opinion in Systems Biology. 2021;28:100401. doi: https://doi.org/10.1016/j.coisb.2021.100401.

10. Piskounova E, Agathocleous M, Murphy MM, Hu Z, Huddlestun SE, Zhao Z, Leitch AM, Johnson TM, DeBerardinis RJ, Morrison SJ. Oxidative stress inhibits distant metastasis by human melanoma cells. Nature. 2015;527(7577):186–91. Epub 2015/10/16. doi: 10.1038/nature15726. PubMed PMID: 26466563; PMCID: PMC4644103.

11. Ubellacker JM, Tasdogan A, Ramesh V, Shen B, Mitchell EC, Martin-Sandoval MS, Gu Z, McCormick ML, Durham AB, Spitz DR, Zhao Z, Mathews TP, Morrison SJ. Lymph protects metastasizing melanoma cells from ferroptosis. Nature. 2020;585(7823):113–8. Epub 2020/08/21. doi: 10.1038/s41586-020-2623-z. PubMed PMID: 32814895; PMCID: PMC7484468.

12. Pascual G, Avgustinova A, Mejetta S, Martin M, Castellanos A, Attolini CS, Berenguer A, Prats N, Toll A, Hueto JA, Bescos C, Di Croce L, Benitah SA. Targeting metastasis-initiating cells through the fatty acid receptor CD36. Nature. 2017;541(7635):41–5. Epub 2016/12/16. doi: 10.1038/nature20791. PubMed PMID: 27974793.

13. Antalis CJ, Uchida A, Buhman KK, Siddiqui RA. Migration of MDA-MB-231 breast cancer cells depends on the availability of exogenous lipids and cholesterol esterification. Clin Exp Metastasis. 2011;28(8):733–41. Epub 2011/07/12. doi: 10.1007/s10585-011-9405-9. PubMed PMID: 21744083.

14. Nath A, Chan C. Genetic alterations in fatty acid transport and metabolism genes are associated with metastatic progression and poor prognosis of human cancers. Sci Rep. 2016;6:18669. Epub 2016/01/05. doi: 10.1038/srep18669. PubMed PMID: 26725848; PMCID: PMC4698658.

15. Ju HQ, Lin JF, Tian T, Xie D, Xu RH. NADPH homeostasis in cancer: functions, mechanisms and therapeutic implications. Signal Transduct Target Ther. 2020;5(1):231. Epub 2020/10/09. doi: 10.1038/s41392-020-00326-0. PubMed PMID: 33028807; PMCID: PMC7542157.

16. Ying W. NAD+/NADH and NADP+/NADPH in cellular functions and cell death: regulation and biological consequences. Antioxid Redox Signal. 2008;10(2):179–206. Epub 2007/11/21. doi: 10.1089/ars.2007.1672. PubMed PMID: 18020963.

17. Cracan V, Titov DV, Shen H, Grabarek Z, Mootha VK. A genetically encoded tool for manipulation of NADP(+)/NADPH in living cells. Nat Chem Biol. 2017;13(10):1088–95. Epub 2017/08/15. doi: 10.1038/nchembio.2454. PubMed PMID: 28805804; PMCID: PMC5605434.

18. Elia I, Broekaert D, Christen S, Boon R, Radaelli E, Orth MF, Verfaillie C, Grünewald TGP, Fendt SM. Proline metabolism supports metastasis formation and could be inhibited to selectively target metastasizing cancer cells. Nat Commun. 2017;8:15267. Epub 2017/05/11. doi: 10.1038/ncomms15267. PubMed PMID: 28492237; PMCID: PMC5437289.

19. Aslakson CJ, Miller FR. Selective events in the metastatic process defined by analysis of the sequential dissemination of subpopulations of a mouse mammary tumor. Cancer Res. 1992;52(6):1399–405. Epub 1992/03/15. PubMed PMID: 1540948.

20. Schild T, McReynolds MR, Shea C, Low V, Schaffer BE, Asara JM, Piskounova E, Dephoure N, Rabinowitz JD, Gomes AP, Blenis J. NADK is activated by oncogenic signaling to sustain pancreatic ductal adenocarcinoma. Cell Rep. 2021;35(11):109238. Epub 2021/06/17. doi: 10.1016/j.celrep.2021.109238. PubMed PMID: 34133937.

21. Minn AJ, Gupta GP, Siegel PM, Bos PD, Shu W, Giri DD, Viale A, Olshen AB, Gerald WL, Massague J. Genes that mediate breast cancer metastasis to lung. Nature. 2005;436(7050):518–24. Epub 2005/07/29. doi: 10.1038/nature03799. PubMed PMID: 16049480; PMCID: 1283098.

22. Padua D, Massague J. Roles of TGFbeta in metastasis. Cell Res. 2009;19(1):89–102. doi: 10.1038/cr.2008.316. PubMed PMID: 19050696.

23. Gomes AP, Ilter D, Low V, Rosenzweig A, Shen ZJ, Schild T, Rivas MA, Er EE, McNally DR, Mutvei AP, Han J, Ou YH, Cavaliere P, Mullarky E, Nagiec M, Shin S, Yoon SO, Dephoure N, Massague J, Melnick AM, Cantley LC, Tyler JK, Blenis J. Dynamic Incorporation of Histone H3 Variants into Chromatin Is Essential for Acquisition of Aggressive Traits and Metastatic Colonization. Cancer Cell. 2019;36(4):402–17 e13. Epub 2019/10/01. doi: 10.1016/j.ccell.2019.08.006. PubMed PMID: 31564638; PMCID: PMC6801101.

24. Ray-Gallet D, Woolfe A, Vassias I, Pellentz C, Lacoste N, Puri A, Schultz DC, Pchelintsev NA, Adams PD, Jansen LE, Almouzni G. Dynamics of histone H3 deposition in vivo reveal a nucleosome gap-filling mechanism for H3.3 to maintain chromatin integrity. Mol Cell. 2011;44(6):928–41. doi: 10.1016/j.molcel.2011.12.006. PubMed PMID: 22195966.

## References

1 Su, X., Lu, W. & Rabinowitz, J. D. Metabolite Spectral Accuracy on Orbitraps. Anal Chem 89, 5940–5948, doi:10.1021/acs.analchem.7b00396 (2017).

2 Cox, A. G. et al. Mitochondrial peroxiredoxin 3 is more resilient to hyperoxidation than cytoplasmic peroxiredoxins. Biochem J 421, 51–58, doi:10.1042/BJ20090242 (2009).

3 Gomes, A. P. et al. Dynamic Incorporation of Histone H3 Variants into Chromatin Is Essential for Acquisition of Aggressive Traits and Metastatic Colonization. Cancer Cell 36, 402–417 e413, doi:10.1016/j.ccell.2019.08.006 (2019).

4 Freese, N. H., Norris, D. C. & Loraine, A. E. Integrated genome browser: visual analytics platform for genomics. Bioinformatics 32, 2089–2095, doi:10.1093/bioinformatics/btw069 (2016).

