## Supplementary figures and images for "NADK upregulation is an essential metabolic adaptation that enables breast cancer metastatic colonization"

### Extended Data Fig. 1

#### Extended Data Figure 1

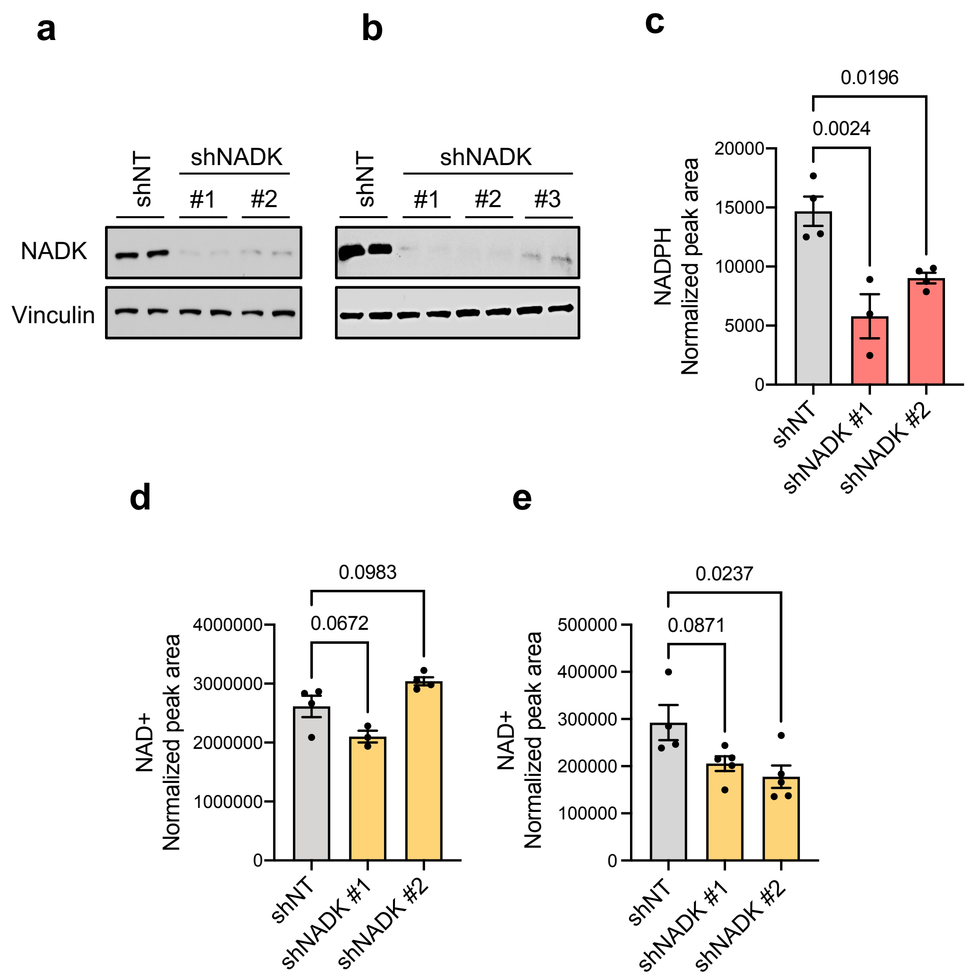

### Extended Data Fig. 2

#### Extended Data Figure 2

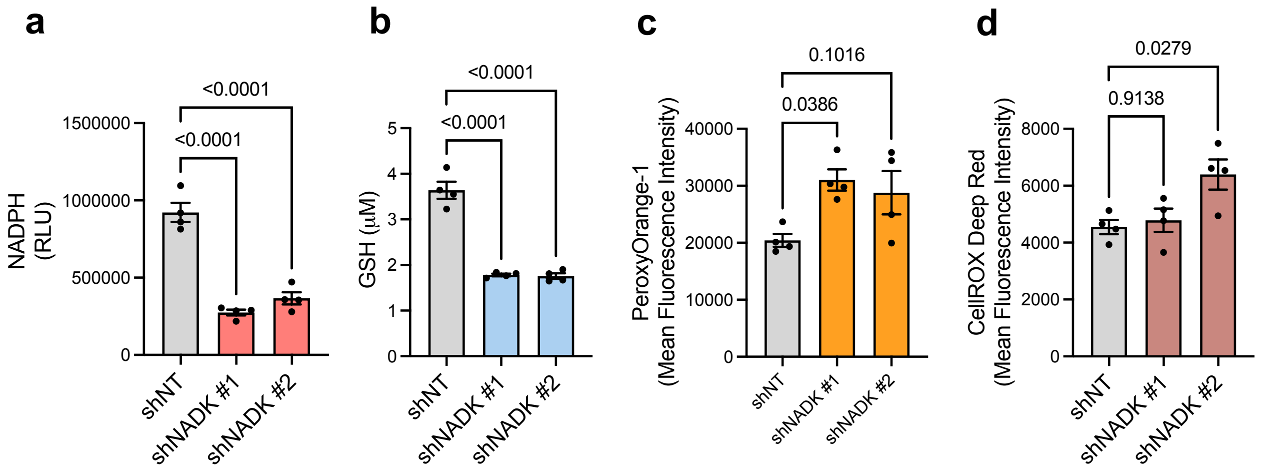


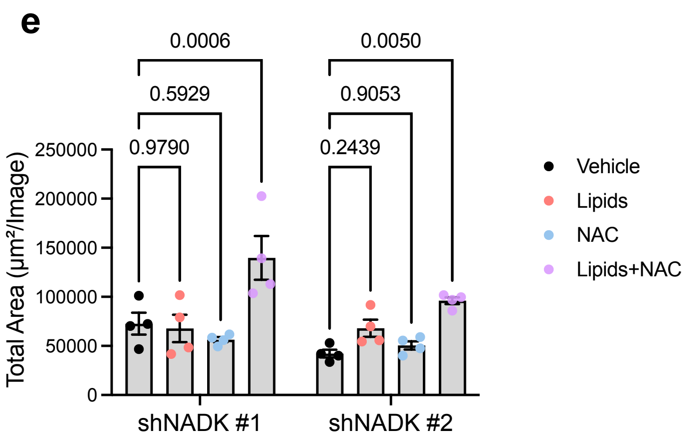

### Extended Data Fig. 3

#### Extended Data Figure 3

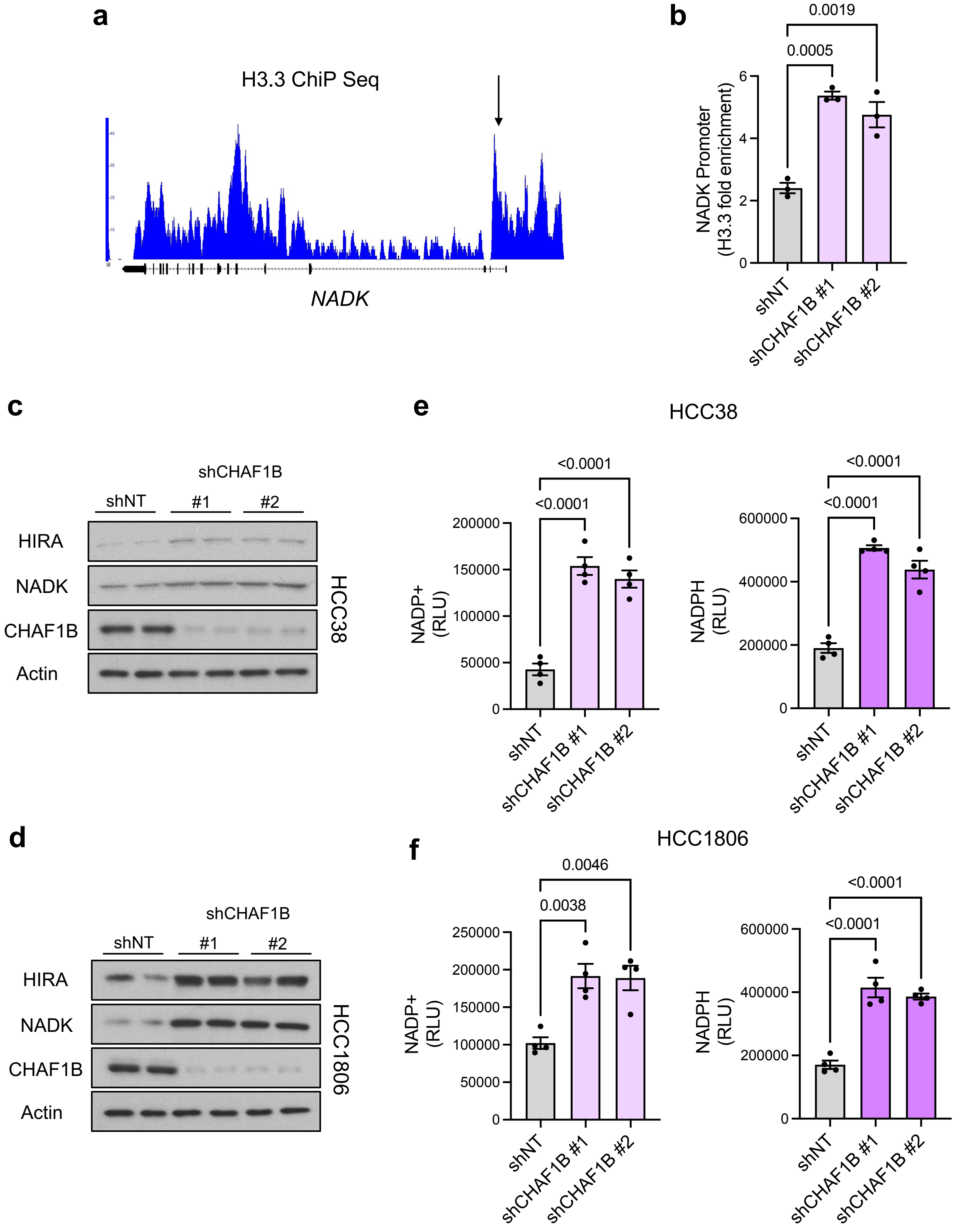

### Extended Data Fig. 4

#### Extended Data Figure 4

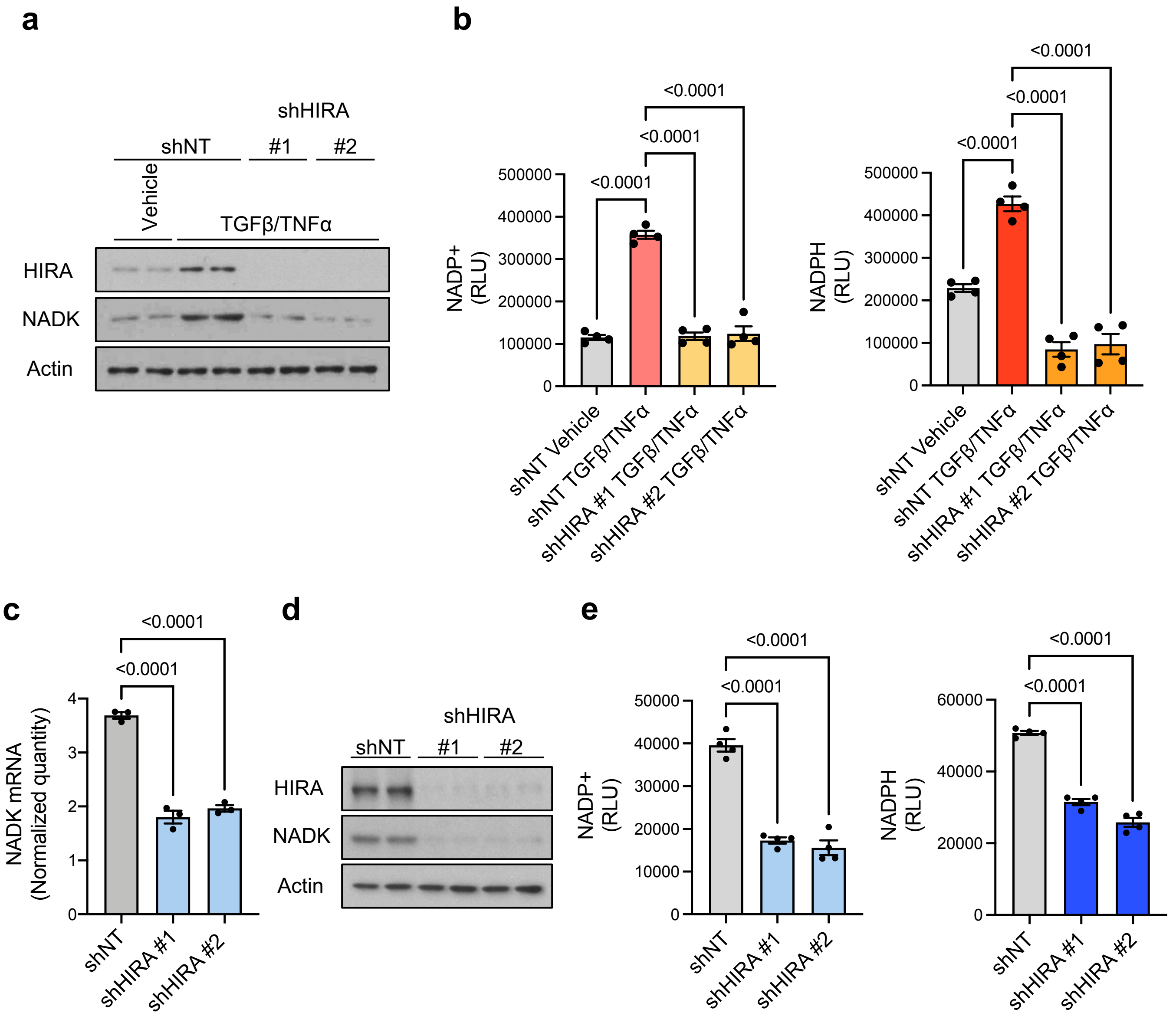
